# Specific Inhibition of GSK-3β by Tideglusib: Potential Therapeutic Target for Neuroblastoma Cancer Stem Cells

**DOI:** 10.1101/2020.02.18.953596

**Authors:** Hisham F. Bahmad, Reda M. Chalhoub, Hayat Harati, Jolie Bou-Gharios, Farah Ballout, Alissar Monzer, Hiba Msheik, Sahar Assi, Tarek Araji, Mohamad K. Elajami, Paola Ghanem, Farah Chamaa, Humam Kadara, Tamara Abou-Antoun, Georges Daoud, Youssef Fares, Wassim Abou-Kheir

**Affiliations:** Department of Anatomy, Cell Biology and Physiological Sciences, Faculty of Medicine, American University of Beirut, Beirut, Lebanon; Neuroscience Research Center, Faculty of Medicine, Lebanese University, Beirut, Lebanon; Department of Translational Molecular Pathology, The University of Texas MD Anderson Cancer Center, Houston, TX, USA; School of Pharmacy, Department of Pharmaceutical Sciences, Lebanese American University, Byblos, Lebanon

**Keywords:** neuroblastoma, GSK-3β, Tideglusib, cancer stem cells, targeted therapy

## Abstract

Neuroblastoma is an embryonic tumor that represents the most common extracranial solid tumor in children. Resistance to therapy is attributed, in part, to the persistence of a subpopulation of slowly dividing cancer stem cells (CSCs) within those tumors. Glycogen synthase kinase (GSK)-3β is an active proline-directed serine/threonine kinase, well-known to be involved in different signaling pathways entangled in the pathophysiology of neuroblastoma. This study aims to assess the potency of an irreversible GSK-3β inhibitor drug, Tideglusib (TDG), in suppressing proliferation, viability, and migration of human neuroblastoma cell lines, as well as its effects on their CSCs subpopulation *in vitro* and *in vivo*. Our results showed that treatment with TDG significantly reduced cell proliferation, viability, and migration of SK-N-SH and SH-SY5Y cells. TDG also significantly inhibited neurospheres formation capability in both cell lines, eradicating the self-renewal ability of highly resistant CSCs. Importantly, TDG potently inhibited neuroblastoma tumor growth and progression *in vivo*. In conclusion, TDG proved to be an effective *in vitro* and *in vivo* treatment for neuroblastoma cell lines and may hence serve as a potential adjuvant therapeutic agent for this aggressive nervous system tumor.

## 1. Introduction

Neuroblastoma (NB) is a common childhood tumor that originates from embryonic neural crest cells that serve as precursor cells of the sympathetic nervous system. It represents the most common extracranial solid tumor among pediatric patients [1], accounting for 6% of all cancer diagnoses of children (0-14 years) in the US [2]. This disease is remarkable for its varied clinical outcomes, whereas some tumors spontaneously regress into mature non-malignant tissue, while others progress and metastasize regardless of intense treatment measures [1, 3]. Current treatment regimens include an induction radiotherapy, surgical excision of the tumor, and high-dose chemotherapy regimen [4]. Even though the overall 5-year survival rate of the NB patients in the US remains around 80% [2], patients with high-risk disease (stage 4, amplified MYCN) – accounting for the majority of the diagnoses – suffer from a long-term survival rate below 50% [5], with cure relapse and tumor recurrence seen in almost 50% of the cases [6].

The malignant recurrence of NB after complete clinical remission, as well as other solid tumors, is notably attributed to the failure in the complete eradication of cancer stem cells (CSC), a subpopulation of cells within the tumor bulk that possess an indefinite self-renewal ability, and play an integral role in tumor initiation, progression, and recurrence [7]. One of the main characteristics of CSC is their potential to resist conventional treatment regimens through the development of multi-drug resistance as seen NB patients and neuroblastoma cell lines *in vitro* [8, 9], which highlights the need to develop effective targeted therapy able to target this subpopulation.

Glycogen Synthase Kinase 3 Beta (GSK-3β) is an active proline-directed serine/threonine kinase that plays a regulatory role in glucose metabolism, as well as other signaling pathways, including cell-fate determination, cellular differentiation and cell division [10]. GSK3-β plays a controversial role in cancer pathophysiology: while it plays a tumor suppressor protein by activating the adenomatous polyposis coli (APC)- β-Catenin destruction complex in colon cancers, recent studies investigated its potential role as a target protein in other cancers, such as pancreatic adenocarcinoma and acute myeloid leukemias [11-14], suggesting non-conventional mechanisms by which GSK-3β regulates carcinogenesis.

In this study, we tested the effect of GSK-3β inhibition by Tideglusib (TDG) on NB cell lines *in vitro* and *in vivo*. Tideglusib is a well-tolerated, irreversible non-ATP competitive GSK-3β inhibitor that was clinically tested for its effect on different neurological disorders, including Alzheimer’s disease and progressive supranuclear palsy [15-17]. Here, we show that treating human NB cell lines with TDG hinders cellular proliferation, decreases cellular survival, and blocks cellular migration. Furthermore, we further focused on its effects on the CSC subpopulation in NB cell lines, using a 3D-sphere formation and propagation model [18-20] over 4 sequential generations of neurospheres.

## 2. Materials and Methods

### 2.1. Cell Culture

Two neuroblastoma cell lines SK-N-SH (ATCC^®^ HTB-11™, USA) [21] and SH-SY5Y (ATCC^®^ CRL-2266™, USA) [22, 23], were cultured and maintained in Dulbecco’s Modified Eagle Media (DMEM) Ham’s F12 (Sigma-Aldrich; cat #D8437), supplemented with 10% of heat inactivated fetal bovine serum (FBS) (Sigma-Aldrich; cat #F9665), 1% Penicillin/Streptomycin (Biowest; cat #L0022-100) and Plasmocin™ prophylactic (Invivogen; cat #ant-mpp). Cell lines were checked using the ICLAC Database of Cross-contaminated or Misidentified Cell Lines confirming they are not misidentified or contaminated. Cells were incubated at 37□ in a humidified incubator containing 5% CO_2_. Tideglusib (TDG) was purchased from Sigma-Aldrich (Cat. # SML0339-10MG; Lot # 123M4615V and 016M4605V) and reconstituted in dimethyl suldoxide (DMSO; Amresco; cat #0231-500ML), per manufacturer’s instructions.

### 2.2. MTT/Cell Proliferation Assay

The anti-proliferative effects of Tideglusib (TDG) on the used cell lines were measured *in vitro* using MTT ([3-(4, 5-dimethylthiazol-2-yl)-2, 5-diphenyltetrazolium bromide]) (Sigma-Aldrich; cat #M5655-1G) assay according to the manufacturer’s instructions [24-26]. In brief, cells (6×10^3^ cells in 100µL full media, per well) were seeded in 3 different 96-well culture plates (one for every time point: 24h, 48h and 72h) and incubated overnight at 37□ and 5% CO_2_. Wells were randomly distributed across different treatment conditions – 3 wells/condition (Control: media only, Vehicle: Media + 0.1% DMSO, treatment groups: 5µM, 25µM, and 50µM TDG in full media). At every time point, media was removed and replaced with fresh media along with 10µL of MTT yellow dye (5mg/mL in DMSO) per well. Afterwards, the cells were incubated for 4 hours, after which 100µL of the solubilizing agent was added to each well. The plates were incubated overnight at room temperature, and the absorbance intensity of every well was measured by the microplate ELISA reader (Multiscan EX) at 595nm. The percentage of cell proliferation was presented as an optical density (OD) ratio of the treated to the untreated cells (control).

### 2.3. Trypan Blue/Cell Viability Assay

The effects of TDG on cell viability was measured *in vitro* using the trypan blue assay [27]. In brief, SH-SY5Y and SK-N-SH cells (60×10^3^ cells/well in 500µL full media) were seeded in 3 different 12-well culture plates (one for each time point: 24h, 48h, and 72h). Cells were incubated overnight at 37□ and 5% CO_2_. Wells were randomly distributed, in duplicate, across different treatment conditions (Control: media only, Vehicle: Media + 0.1% DMSO, treatment groups: 5µM, 25µM, and 50µM TDG in full media). At every time point, cells from each well were treated with Trypsin and viable cells were counted on a hemocytometer under an inverted light microscope after staining cell suspension with Trypan blue. The percentage of cell viability was determined as a ratio of viable cells counted in treated to untreated conditions.

### 2.4. Wound Healing Assay

Wound healing assay was used to assess the effects of TDG on cell migration. SH-SY5Y and SK-N-SH cells (5×10^5^ cells/well) were seeded in 6-well culture plates and incubated at 37□ and 5% CO_2_ until they reached 90% confluence. Cells were then treated with 5mg/mL of Mitomycin C (Sigma-Aldrich; cat #M0503-5×2MG) for 1 hour to block cellular proliferation. A sterile 200µL pipet was used to create two scratches per well in each monolayer. Cells were then washed twice with dulbecco’s phosphate buffered saline (D-PBS) (Sigma-Aldrich; cat #D8537-500ML) to remove cell debris. The remaining cells were distributed into three conditions: Control (Full media), Vehicle (full media + 0.1% DMSO), and treatment (25µM TDG in full media). Pictures of the scratches were taken using an inverted light microscope at the following time points: 0h, 6h, 12h, 24h, and 48h. The distance travelled by the cells was measured using Zen Microscope Software (Zen 2.3).

### 2.5. 3D culture and Sphere-Formation Assay

The sphere formation assay was used as previously reported by our lab [19, 20]. In brief, single SH-SY5Y and SK-N-SH cell suspensions (10×10^3^ cells/well) were seeded in Matrigel™/serum free DMEM Ham’s F-12 (1:1). The solution was then plated gently around the rim of individual wells of 24-well culture plate (50µL per well). The Matrigel™ (Corning Life Sciences; cat #354230) was allowed to solidify for 1 hour at 37°C in a humidified incubator. Wells were randomly assigned to control and treatment conditions (control, 0.1µM, 1µM, and 5µM TDG). 500 µL/well of complete media (+5% FBS) was gently added to the center of each well and changed regularly every 3 days. At day 9 after plating, spheres were pictured and counted. SH-SY5Y spheres were further harvested for propagation.

For spheres propagation, the medium was aspirated from the center of the wells. The Matrigel™ was digested with 0.5mL of 1mg/mL Dispase II solution (ThermoFisher; cat #17105-041) dissolved in serum-free DMEM Ham’s F-12 for 60 minutes at 37°C in a humidified incubator. SH-SY5Y and SK-N-SH spheres were collected and incubated in warm trypsin at 37°C for 5 minutes; trypsin was used to dissociate spheres into single cell suspensions. Cells were counted and re-seeded at 2×10^3^ cells/well. The propagation of the spheres was repeated over 4 generations. The sphere forming unit (SFU) was calculated as the ratio of the number of spheres counted at day 9 to the number of cells originally seeded. Bright field images of the spheres were obtained using Axiovert microscope from Zeiss at 5× magnification.

### 2.6. Western Blotting Analysis

For 2D, SH-SY5Y cells were cultured in 6-well plates (5×10^5^ cells/well) until they reached 70% confluency. Three wells were then treated with 25µM TDG for 48 hours, while the remaining wells were taken as control. The plates were then washed with ice-cold D-PBS to remove any residual media. Similarly, for 3D, SH-SY5Y spheres (G1) were treated with 5µM TDG while others were taken as control. G1 spheres were collected and washed with ice-cold D-PBS. Cells/spheres were treated and lysed using radioimmunoprecipitation (RIPA) buffer (0.1% sodium dodecyl sulfate (SDS) (v/v), 0.5% sodium deoxylate (v/v), 150mM sodium chloride (NaCl), 100mM EDTA, 50mM Tris-HCl (pH=8), 1% Tergitol (NP40) (v/v), 1mM PMSF, and protease and phosphatase inhibitors (one tablet of each in 10mL buffer, Roche, Germany)), scraped off the plates, transferred into micro-centrifuge tubes and incubated on ice for 30 minutes. Sonication was used to maximize the protein yield. Lysates were then centrifuged at 13,600 rpm for 15 minutes at 4°C, to pellet the cell debris.

Protein concentrations of the collected supernatants were quantified using DC™ Protein Assay (Bio-Rad). For immunoblotting, 50μg of proteins were electrophoresed in 8% or 12% polyacrylamide gel and then transferred to PVDF membranes (Bio-Rad Laboratory, CA, USA) overnight. Membranes were blocked with 5% bovine serum albumin (BSA) (v/v) (Amresco; cat #0332-100G) for 2 hours and blotted at 4°C overnight with primary antibodies as follows: rabbit anti-phospho-GSK-3β (Ser9) (5B3) (1/500 dilution; Cell Signaling; cat #9323), rabbit anti-GSK-3β (1/1000 dilution; Cell Signaling; cat #27C10,), and mouse anti-GAPDH (1/5000 dilution; Novus Biologicals; cat #NB300-221). The next day, membranes were washed and incubated at room temperature for 2 hours with the appropriate HRP-conjugated secondary antibodies as follows: mouse anti-rabbit (1/1000 dilution; Santa Cruz; cat #sc-2357) and mouse IgGκ BP (1/1000 dilution; Santa Cruz; cat #sc-516102). Finally, bands were detected using Lumi-Light Western Blotting Substrate (Roche; cat #12015200001) and visualized using autoradiography. Band intensities were digitized and analyzed using ImageJ software.

### 2.7. Mouse Neuroblastoma Xenografts

This study was approved by the Institutional Animal Care and Utilization Committee (IACUC) of the American University of Beirut. Neuroblastoma xenografts were generated using mouse SH-SY5Y cells. Cells were injected at a concentration of 1.2×10^6^ cells in 100μL total volume of cells and Matrigel^TM^ (1:1) using a 27 G needle subcutaneously, into the flanks of NOD-SCID male mice (6-8 weeks old) [28]. Once palpable tumor (approximate size 1mm^3^) was detected, mice were intraperitoneally injected 3 times a week for 2 weeks with 20mg/kg TDG or vehicle only (Lipofundin/DMSO) and tumor volumes were measured every 3 days by direct physical measurements using a digital caliper (Model DC150-S) to determine tumor size and expansion. Mice weight was monitored at the initiation of the experiment and at the time of sacrifice. The following formula for volume assessment was applied: V = (3.14/6)×L×W×H; where V is the tumor volume in mm^3^, L is the tumor length in mm, W is the tumor width in mm, and H is the tumor height in mm. Measurements were carried out until the termination of the experiment. Data represent an average of n=3 mice. The data are reported as mean ± SEM.

### 2.8. Data Analyses

Statistical analysis was performed using GraphPad Prism 7 software. The significance of the data was determined using proper statistical tests, including the student *t*-test and the two-way ANOVA statistical test, followed by multiple comparisons using Bonferroni post-hoc analysis. P-values of *p<0*.*05* (*), *p<0*.*01* (**) and *p<0*.*001* (***) were labeled significant, highly significant and very highly significant, respectively.

## 3. Results

### 3.1. GSK-3β mRNA Expression Patterns in Human Neuroblastoma Tissues

In our study, we first aimed to assess the expression pattern of *GSK-3β* gene in human neuroblastoma tumor tissues as compared to other body cancer tissues. For this, we surveyed a publicly available dataset (Neale Multi-cancer Statistics, 60 samples; data retrieved from Oncomine.org) encompassing human tumor tissues from different organs. mRNA expression analysis revealed high expression of *GSK-3β* gene among neuroblastoma tissues relative to other organ specific tumor tissues in three out of four probes of the dataset (**Fold change = 1.639; *p = 4*.*06E-4***) (**Fig 1 and Supp Fig 1**).

**Fig 1.**
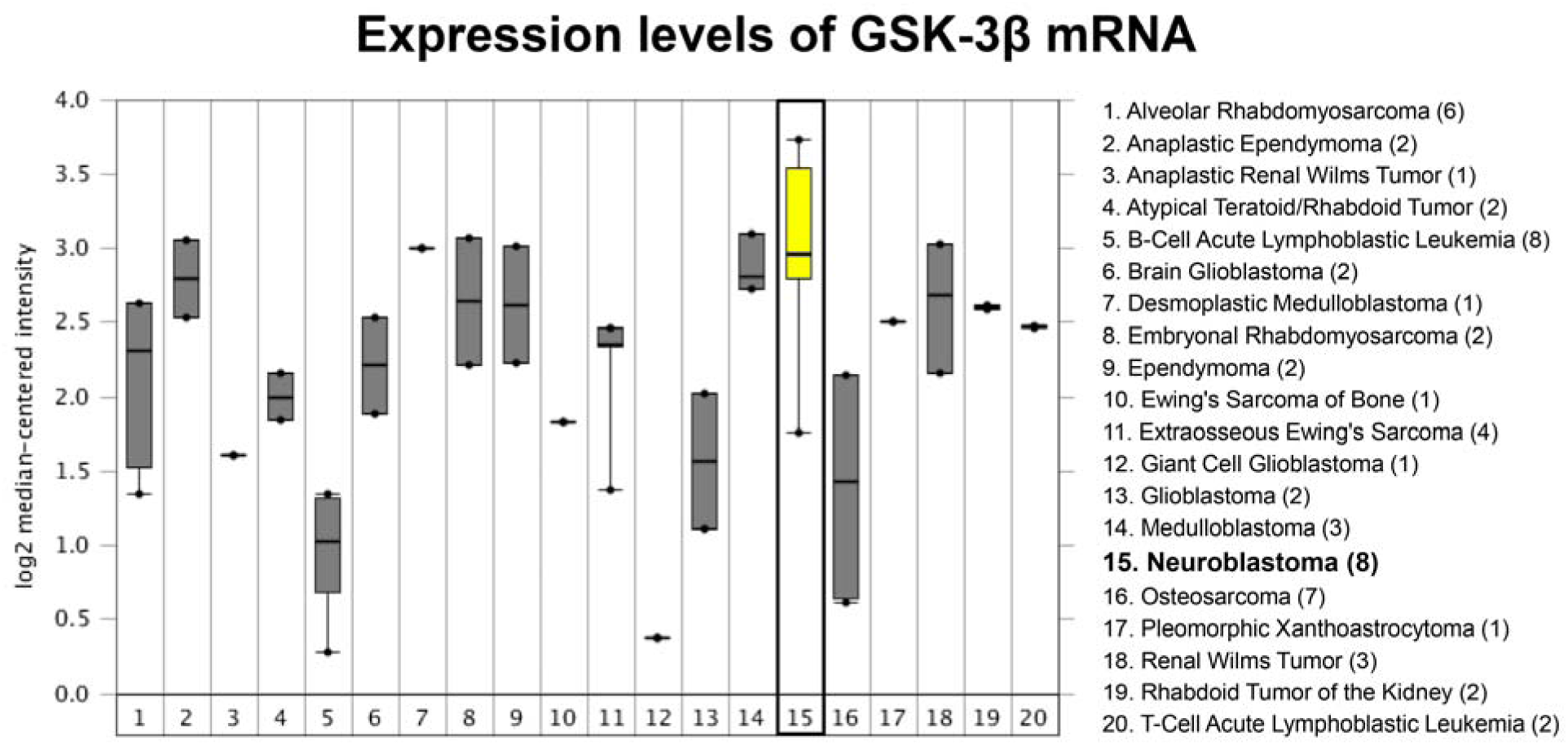
Expression levels of *GSK-3β* mRNA were assessed in an array set comprised of human pan-tumor samples (Neale Multi-cancer Statistics, Reporter 226183_at is presented; the remaining probes of the dataset are presented in Supp Fig 1: Reporters 209945_s_at, 226191_at, and 242336_at). Expression within tumor tissues was presented by log (base 2) median-centered expression of *GSK-3β*. Box and whiskers plots indicate median and interquartile range. *p* values were obtained using *t*-tests (Neale Multi-cancer Statistics, 60 samples; data retrieved from Oncomine.org). Analysis revealed that mRNA expression of *GSK-3β* gene was the highest among neuroblastoma tissues relative to other organ specific tumor tissues (Fold change = 1.639; *p = 4*.*06E-4*).

### 3.2. Tideglusib decreases cell viability and cell proliferation of human NB cell lines

The effect of TDG on the cellular proliferation and cellular viability of human neuroblastoma cell lines, SK-N-SH and SH-SY5Y, was assessed *in vitro* using the MTT Assay (**Fig 2A and 2B**) and the Trypan Blue assay, respectively (**Fig 2C and 2D**). TDG significantly inhibited the proliferative ability of SH-SY5Y and SK-N-SH cell lines in a dose-dependent manner (two-way ANOVA showed significant effect for treatment *p<0*.*0001*, for both cell lines). TDG treatment of 25µM achieved nearly a 50% inhibitory effect on both cell lines, after 72h (**Fig 2A and 2B**). In addition, for further validation, we saw a significant effect of TDG treatment on cell viability using the trypan blue exclusion assay. At 72h of treatment with 25µM of TDG, there was a significant decrease in the number of viable cells in culture, for both SH-SY5Y and SK-N-SH cell lines (**Fig 2C and 2D**).

**Fig 2.**
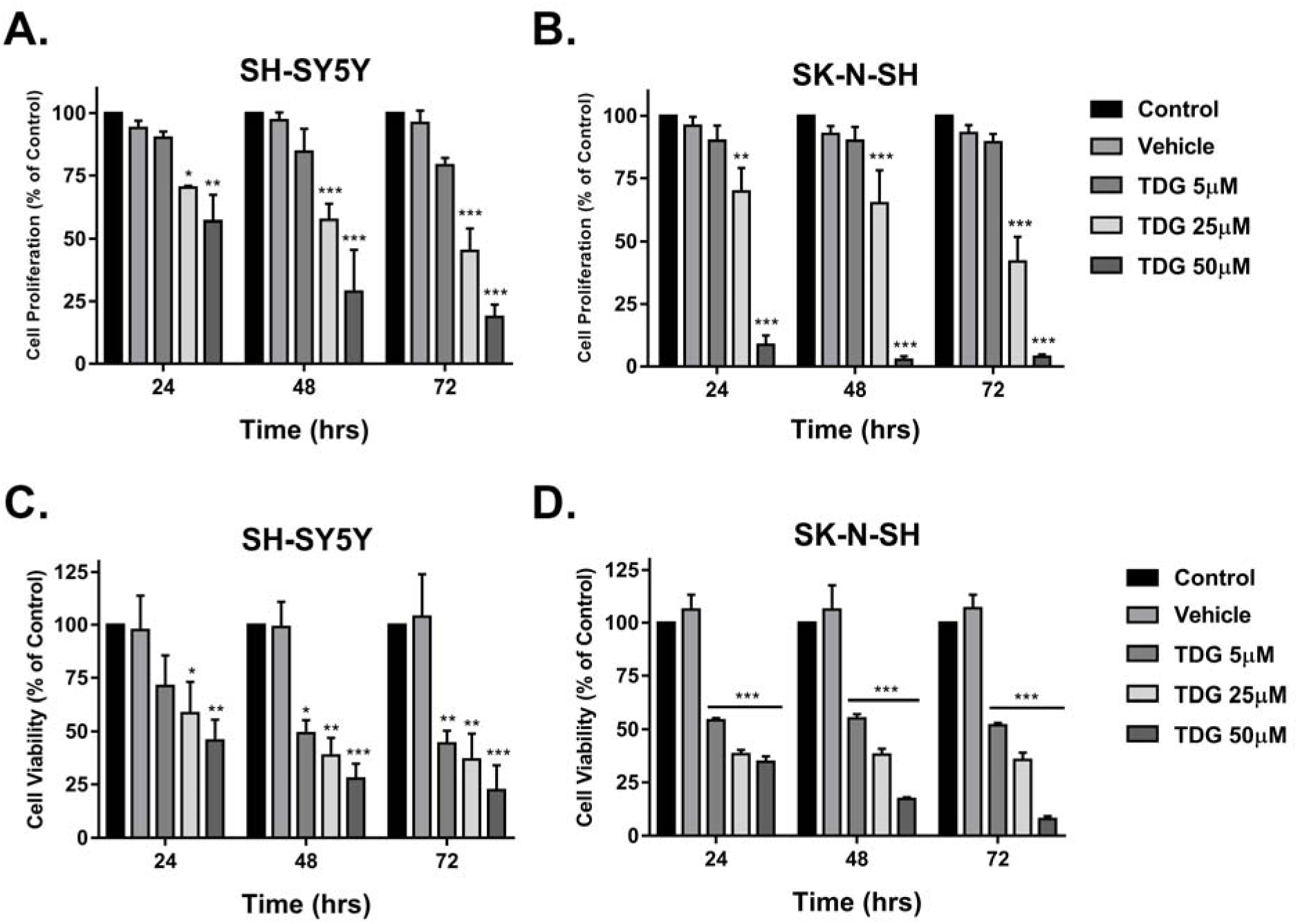
Tideglusib significantly decreases cell proliferation and cell viability of human neuroblastoma cells. **(A)** The effect of TDG on cell proliferation was determined using the MTT assay. Tideglusib significantly decreases cell proliferation of SK-N-SH (two-way ANOVA; treatment F_(4, 30)_ = 143, *p<0*.*001*; time F_(2, 30)_ = 2.02, *p=0*.*15*; interaction F_(8, 30)_ = 1.29, *p=0*.*2858*) and SH-SY5Y (two-way ANOVA; treatment F_(4, 30)_ = 50.78, *p<0*.*001*; time F_(2, 30)_ = 5.801, *p=0*.*0074*; interaction F_(8, 30)_ = 1.738, *p=0*.*1303*) cells in dose-dependent manner, as determined by MTT. **(B)** The effect of TDG on cell viability was determined using the trypan blue assay. Tideglusib significantly decreases the percentage of viable cells in SK-N-SH (two-way ANOVA; treatment F_(4, 30)_ = 248.5, *p<0*.*001*; time F_(2, 30)_ = 2.791, *p=0*.*0773*; interaction F_(8,_ 30) = 2.002, *p=0*.*0808*) and SH-SY5Y (two-way ANOVA; treatment F_(4, 30)_ = 25.22, *p<0*.*001*; time F_(2, 30)_ = 2.16, *p=0*.*1329*; interaction F_(8, 30)_ = 0.524, *p=0*.*8289*) cells in dose-dependent manner, as determined by MTT. *The data are reported as mean ± SEM of three independent experiments. Bonferroni post-hoc analysis was done to determine simple factor effects. (**p<0*.*01, ***p<0*.*001)*.

### 3.3. Tideglusib inhibits cell migration of human NB cells *in vitro*

Following that, we assessed the effect of TDG on cellular migration, the main feature that underlie cancer spread and metastasis. This was done using a wound healing assay on both cell lines. Mitomycin C was used to block cell proliferation. In untreated conditions, both cell lines were able to migrate through and close the wounds within 48 hours. Under 25µM treatment with TDG, the wound made in SH-SY5Y and SK-N-SH monolayers remained patent by 60% and 70% respectively (**Fig 3**). This shows that TDG treatment is effective in impeding the migrative ability of neuroblastoma cell lines in culture.

**Fig 3.**
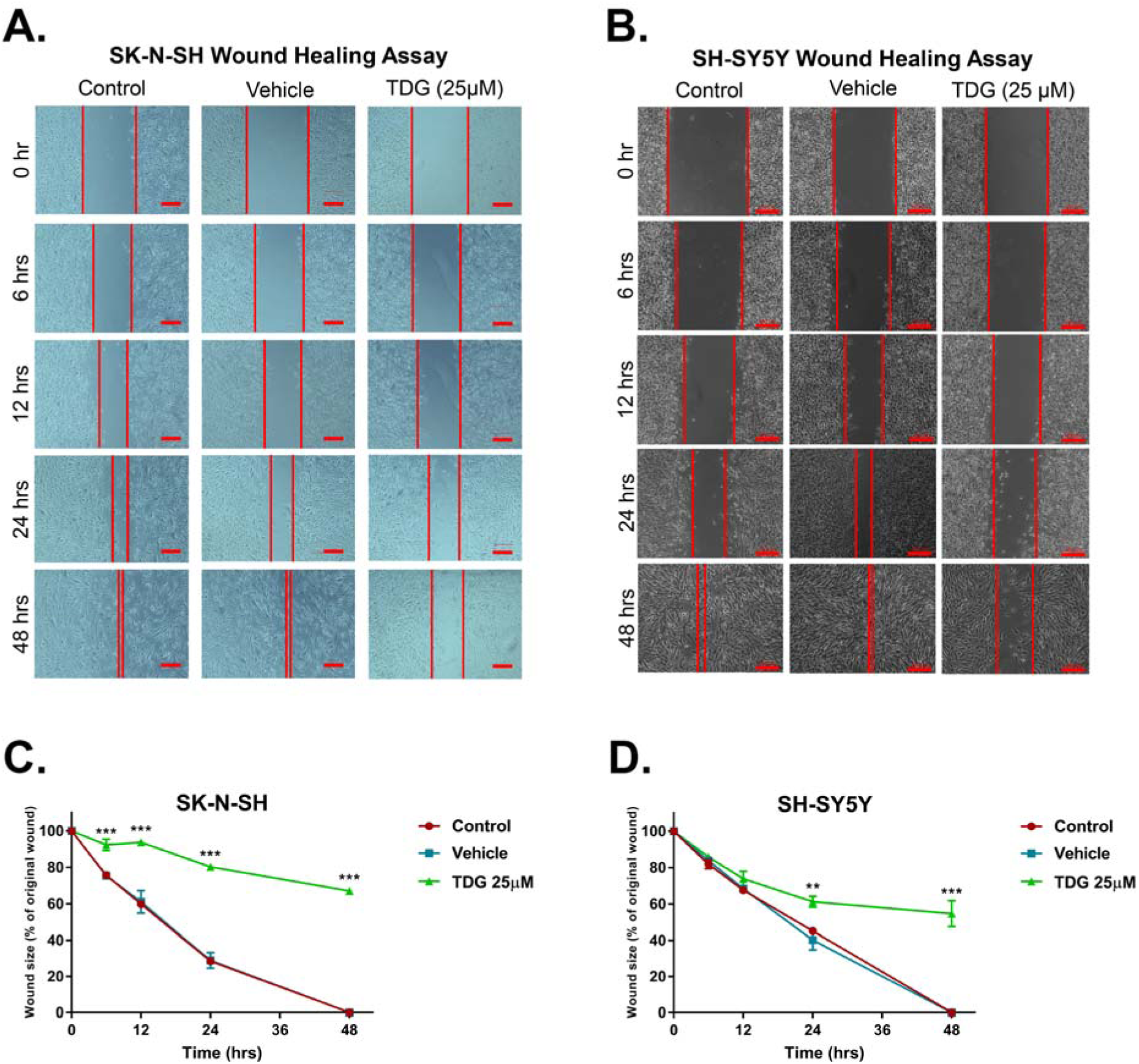
Tideglusib inhibits cell migration of SK-N-SH and SH-SY5Y human neuroblastoma cells. Representative figures showing the scratch made in SK-N-SH **(A)** and SH-SY5Y **(B)** cell lines at five different timepoints: 0h, 6h, 12h, 24h and 48h. These figures show closure of the wound after 48 hours in control and vehicle treated conditions as opposed to the TDG (25 µM) treated conditions. Scale = 200µm in (A) and 100µm in (B). A scratch was made to cells seeded in 6-well plates at T=0h using a 200µL pipet tip; distances between cells were assessed at the different timepoints to determine the drug’s effects on cellular migration. The data are reported as percentages of the distance between cells relative to original wound size at T=0h. Tideglusib (25 µM) significantly inhibited cell migration of SK-N-SH **(C)** (two-way ANOVA with repeated measures: treatment F_(2, 6)_ = 1659, *p<0*.*001*; time F_(4, 24)_ = 454.3, *p<0*.*001*; interaction F_(8, 24)_ = 38.83, *p<0*.*001*) and SH-SY5Y **(D)** (two-way ANOVA with repeated measures: treatment F_(2, 6)_= 80.98, *p<0*.*001*; time F_(4, 24)_ = 340.8, *p<0*.*001*; interaction F_(8, 24)_ = 19.29, *p<0*.*001*) cell lines, in a dose- and time-dependent manners. *The data are reported as mean ± SEM of three independent experiments. Bonferroni post-hoc analysis was done to determine simple factor effects. (**p<0*.*01, ***p<0*.*001 when compared to control)*.

### 3.4. Tideglusib reduces the sphere-forming ability of SH-SY5Y and SK-N-SH cells

Single cell suspensions of SH-SY5Y were cultured under non-adherent conditions in Matrigel™ for 14 days. Sphere forming ability was monitored daily using an inverted light microscope, and pictures were taken to keep track of the size and shape of neurospheres. The sphere formation assay was used as a functional assay to study the stem/progenitor cells subpopulation within SH-SY5Y cell line. Treating cells with TDG after seeding the cells in Matrigel™ significantly decreased the percentage of SFUs in a dose dependent manner (one-way ANOVA, *p=0*.*0037*) (**Fig 4A and 4B**), as well as the average sphere volume (one-way ANOVA, *p<0*.*001*) (**Fig 4C**). Notably, inhibitory effects of TDG were achieved at lower concentration in 3D assay compared to functional assays on cellular monolayers. To further validate our results, we performed the spheres formation assay on SK-N-SH cell lines. Effect of TDG was consistent with that observed with SH-SY5Y where a decrease in the percentage of SFUs at G1 spheres of SK-N-SH was observed in a dose dependent manner (one-way ANOVA, *p<0*.*001*) (**Supp Fig 2A**).

**Fig 4.**
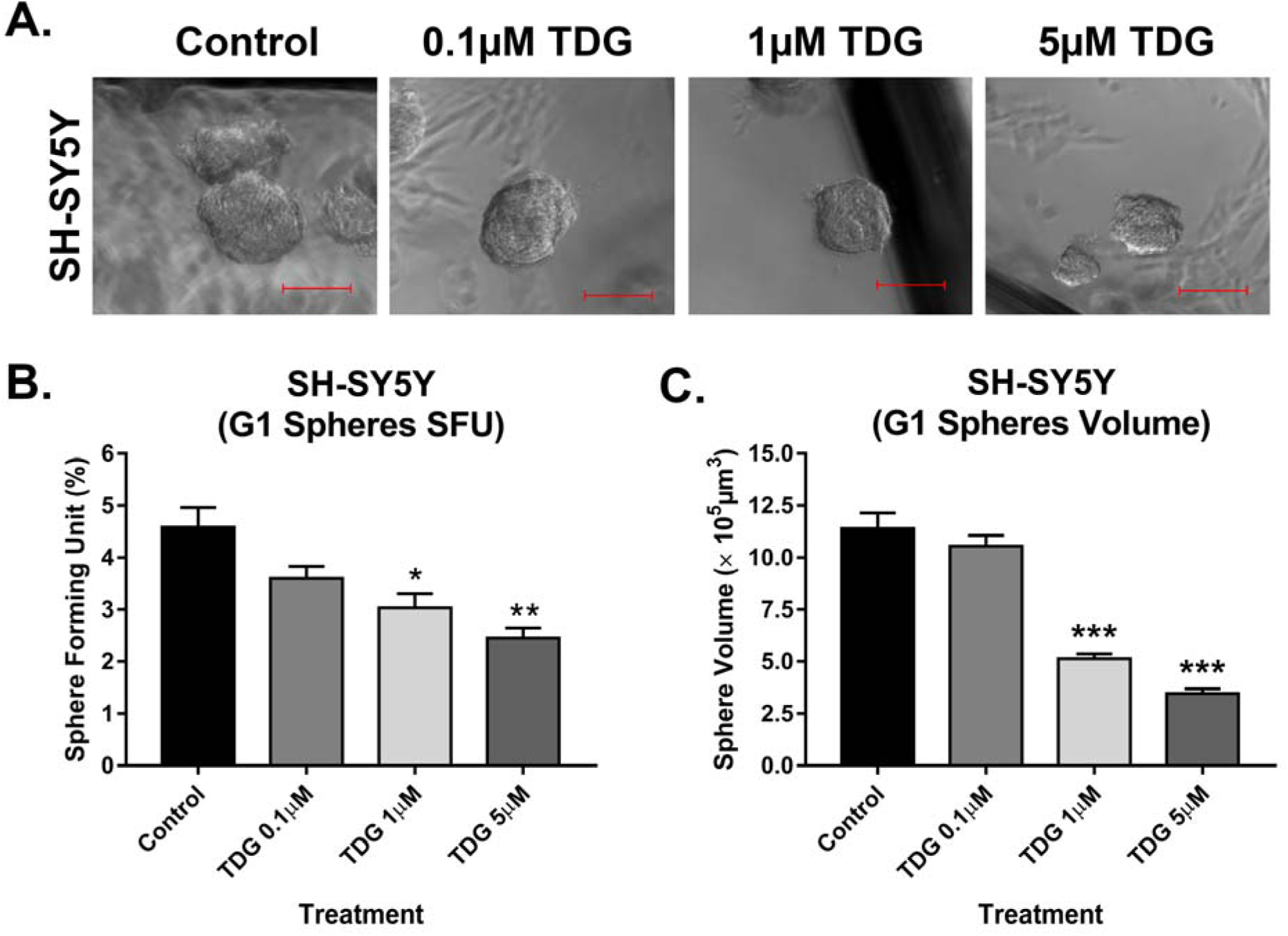
Tideglusib effectively decreases the percentage of self-forming units and volume of spheres in the sphere formation assay on SH-SY5Y cells. **(A)** Representative images taken of SH-SY5Y spheres under different conditions (control, 0.1µM TDG, 1µM TDG and 5µM TDG) using inverted light microscopy showing the gradual decrease in size of spheres in treatment dose-dependent manner. **(B)** Tideglusib decreases the percentage of SFUs in SH-SY5Y cell suspensions in a dose-dependent manner. (One-way ANOVA followed by Bonferroni multiple comparisons: treatment F_(3, 8)_ = 10.6, *p=0*.*0037)* **(C)** TDG treatment decreases the volume of the formed spheres in a dose-dependent manner (one-way ANOVA followed by Bonferroni multiple comparisons: treatment F_(3, 356)_ = 66.27, *p<0*.*001). The data are reported as mean ± SEM of three independent experiments. Bonferroni post-hoc analysis was done to determine simple factor effects. (*p<0*.*05, **p<0*.*01 and ***p<0*.*001 when compared to control)*.

### 3.5. Tideglusib inhibits the sphere self-renewal ability by targeting an enriched population of SH-SY5Y and SK-N-SH cancer stem/progenitor cells

One of the main characteristics of CSCs is their self-renewal ability, largely responsible of cancer recurrence. To study the effect of TDG on this characteristic, we propagated SH-SY5Y and SK-N-SH spheres over multiple generations, wherein the cells taken from one generation of spheres were isolated into single cell suspensions and seeded again under non-adherent conditions. Consecutive generational propagations of those spheres are thought to enrich the cancer stem/progenitor cells subpopulation, by emphasis on their ability of anchorage-independent growth [29]. The experimental design and results of three independent experiments are shown in **Fig 5** for SH-SY5Y and in **Supp Fig 2B** for SK-N-SH. Noteworthy, treating the G4 spheres, which acquired an enriched stem/progenitor subpopulation of cells, with 5µM of TDG significantly decreased the percentage of SFUs by around 95% for SH-SY5Y cells (student independent *t*-test, *p<0*.*001*, **Supp Fig 3A**) and 80% for SK-N-SH cells (student independent *t*-test, *p<0*.*001*, **Supp Fig 3B**).

**Fig 5.**
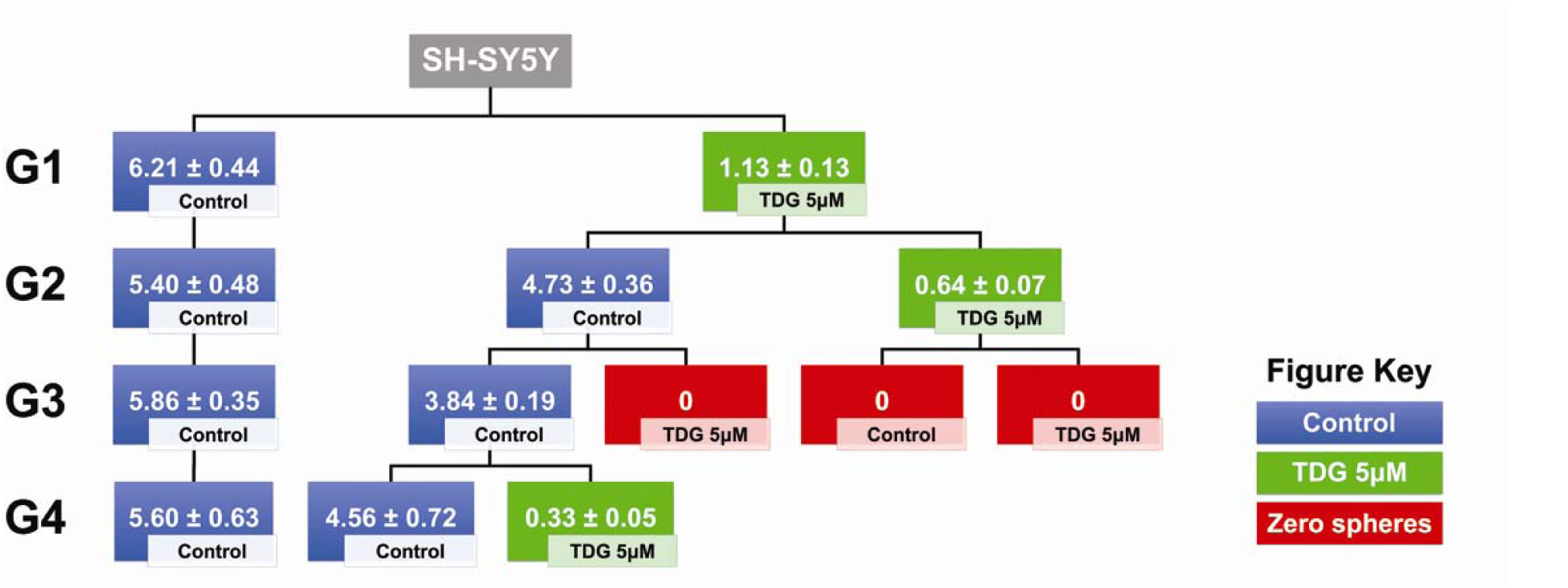
Tideglusib targets an enriched cancer stem/progenitor subpopulation within SH-SY5Y cell line, decreasing SFU across multiple generations. Schematic summarizing the experimental design and results of serial propagation of spheres across 4 generations. Spheres from control and treated conditions were isolated, dissociated into single cell suspensions and seeded under non-adherent conditions. Wells were then randomly distributed into treated and non-treated conditions to assess the effect of treatment across generations. The numbers shown represent the average percentage of SFUs as obtained from three independent experiments. The data was analyzed using multiple independent *t*-tests across each generation.

For SH-SY5Y cells, we decided to test the self-renewal ability of the cells by propagating the same spheres, into two new conditions: control and treated. We noticed that after a single exposure to treated conditions at G1, the SFU significantly dropped to 1.13% compared to 6.21% in control conditions (student independent *t*-test, *p<0*.*0001*) (**Fig 5**). However, once propagated into normal conditions again, the cells successfully regain their self-renewal ability (SFU = 4.73% at G2). According to our data, it takes two treatment regimens in two generations to completely abolish the self-renewal ability of the spheres, i.e. single cell suspensions from spheres previously treated in two generations fail to form any more spheres after propagation (**Fig 5**).

### 3.6. Tideglusib inhibits GSK-3β at protein levels

To validate the direct effect of TDG on its respective target GSK-3β, we used western blotting in order to detect differences in protein expression between the cellular lysates of treated (25µM of TDG) and non-treated SH-SY5Y cells and G1 spheres. GSK-3β inhibition by TDG was established by monitoring the levels of expression of the inhibited form of GSK-3β, phosphorylated at Serine 9 (p-GSK-3β Ser 9). Treating cells with TDG significantly increased the expression of p-GSK-3β (Ser 9) in SH-SY5Y cell lines by around 2.62 times (*p=0*.*0102*) as compared to the control group (**Supp Fig 4A**), signifying GSK-3β inhibition. Besides, treating SH-SY5Y G1 spheres with TDG significantly increased the expression of p-GSK-3β (Ser 9) by around 1.15 times (*p=0*.*0445*) as compared to the control group (**Supp Fig 4B**).

### 3.7. Tideglusib inhibits neuroblastoma growth *in vivo*

Lastly, we assessed the potential effect of TDG on neuroblastoma tumor growth *in vivo*. We injected NOD□SCID mice with SH-SY5Y cells subcutaneously generating neuroblastoma xenografts. The average weight of the mice was monitored throughout the experiment and was maintained within a normal range during the study, signifying that TDG treatment was well tolerated (**Fig 6A**). Treatment with TDG (20mg/kg TDG) resulted in significant inhibition of tumor growth in SH-SY5Y injected mice which was reflected by the reduction in the volume of tumors after 15 days of treatment (**Fig 6B**). These results indicate that TDG significantly reduces neuroblastoma tumor cell growth in xenograft mouse models.

**Fig 6.**
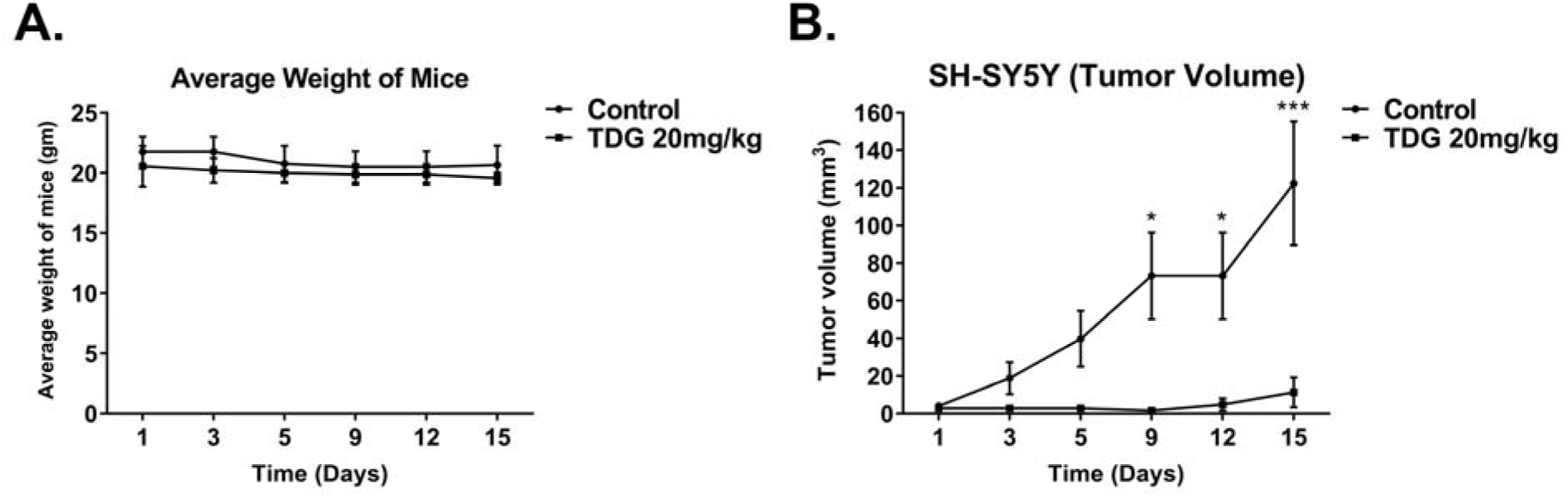
TDG treatment drastically reduces neuroblastoma tumor burden in xenograft mouse models. 1.2□×□10^6^ SH-SY5Y cells were subcutaneously transplanted in 6□ to 8□weeks□old NOD□SCID mice. **(A)** Average weight of mice throughout the experiment was recorded. **(B)** Tumor size measurements were initiated upon the detection of a palpable tumor post□cell injection. Tumor volume was assessed by direct physical measurements of the tumors at the primary site of injection, every 3 days, until the termination of the experiment. The following general formula was applied: *V = (3*.*14/6)×L×W×H)*; where V is the tumor volume in mm^3^, L is the tumor length in mm, W is the tumor width in mm, and H is the tumor height in mm. (two-way ANOVA; treatment F_(1, 24)_ = 37.12, *p<0*.*001*; time F_(5, 24)_ = 5.053, *p=0*.*0026*; interaction F_(5,_ 24) = 3.953, *p=0*.*0093*). Data represent an average of n=3 mice. The data are reported as mean□±□SEM. *(*P<0*.*05; ***P<0*.*001)*.

## 4. Discussion

Neuroblastoma is a solid tumor of the peripheral nervous system that arises from neural crest cells and is typically localized within the medulla of suprarenal glands or within the sympathetic nerve ganglia [3]. The standard care of treatment for most nervous system tumors comprises of maximal surgical resection, radiation therapy, and chemotherapy; yet, tumors eventually recur in the majority of patients despite multimodal treatment. The main reason behind the failure of conventional chemotherapy is hypothesized to be the presence of dormant slowly dividing CSCs within the tumor bulk that develop multi-drug resistance and drive tumor recurrence [30]. Thus, there is ultimate need to come up with novel effective therapies that uniquely target the CSCs population and its related molecular pathways [31]. Several nervous system cancers have been reported to harbor CSCs, such as medulloblastomas [32], glioblastomas [33], and neuroblastomas [20, 34].

Molecular alteration in different signaling pathways of CSCs have been linked to abnormal proliferation, self-renewal and differentiation of these cells, and accordingly many of those pathways have served as potential therapeutic targets and prognostic factors in human oncology [33]. In neuroblastoma tumors [35], some of the oncogenic signaling pathways implicated include PI3K/Akt/mTOR/S6K1, Ras/MAPK, VEGF, EGFR, and p53 [36-38]. Interestingly, GSK-3β represents a signaling node at the crossroads of many of those pathways [12, 20, 39]. This molecule has been associated with several pathological processes in the human body such as bipolar depression, Alzheimer’s disease, Parkinson’s disease and non-insulin-dependent diabetes mellitus (NIDDM) [40].

In oncology, GSK-3β has shown to express opposite actions in different tumors; it has been an oncogenic molecule in some tumors, but a tumor suppressor in others [41]. The exact underlying mechanism of action, at cellular and molecular levels, of GSK-3β in the context of boosting tumor progression is not fully understood; yet, it has been related to blockade of GSK-3β-mediated upregulation of NF-κB-mediated gene transcription [14]. Hence, we hypothesized that targeting this molecule with selective inhibitors, such as TDG - which is now under Phase II Clinical Trials for Alzheimer’s Disease and patients with progressive supranuclear palsy, with minimal adverse effects being reported among patients under study - might carry hope as a novel potential CSCs-targeted therapy to patients suffering from neuroblastoma tumor [15, 42].

First, we surveyed an online publicly available dataset (Neale Multi-cancer Statistics) to determine and compare between mRNA expression patterns of *GSK-3β* in human neuroblastoma tissues and other body tumor tissues. Interestingly, *GSK-3β* was significantly overexpressed in neuroblastoma tissues relative to other tumor tissues, with a fold change of 1.639 (*p = 4*.*06E-4)*.

Next, we assessed the anti-tumor properties and mechanism of action of TDG - an in-clinical-trial drug that specifically inhibits GSK-3β - on two human neuroblastoma cells, SH-SY5Y and SK-N-SH, respectively, and investigated its effect on cell proliferation, viability, and migration *in vitro*, all hallmarks of tumorigenesis. Our results revealed that TDG significantly inhibited the proliferation and survival of both cell lines, in a dose- and time-dependent manners. TDG also significantly reduced migratory ability of both cells. Our results are consistent with those of Mathuram *et al*., where they have shown that TDG, at molecular level, induces apoptosis in human neuroblastoma IMR32 cells, provoking sub-G0/G1 accumulation and ROS generation [43].

Eventually, we also sought to determine the ability of TDG to target the sub-population of cancer stem/progenitor cells in SH-SY5Y cells using a 3D neurospheres formation assay in Matrigel™ *in vitro* [20, 44]. Treatment with TDG at a concentration as low as 0.1µM significantly inhibited SFU as well as sphere size of SH-SY5Y cells. Notably, significant progressive decrease in the number and size of cultured G1 neurospheres followed a dose-dependent manner. Furthermore, consecutive treatment of SH-SY5Y neurospheres at G1 and G2 with the same concentrations of TDG, caused prominent reduction in SFU where neurospheres formation was completely abolished at G3. Sphere formation assay was performed on SK-N-SH cells as well to validate the effect of TDG showing consistent results. Thus, we concluded that TDG is effective in targeting the self-renewal ability of CSCs, a hall mark of cancer progression. When compared to 2D culture, TDG treatment was more potent when used in a 3D culture, whereby lower drug dosages were sufficient to exert an even stronger cytotoxic effect on neuroblastoma cells. Consistent with the *in vitro* data, SH-SY5Y cells treated with TDG *in vivo*, drastically reduced the tumorigenic potential of tumor cells.

Lastly, we believe that our study has some limitations related to the methodology and experimental design. First, we assessed in our work the inhibitory effect of TDG on two human neuroblastoma cell lines as models for this nervous system tumor. Future experiments will follow after acquiring more human cell line models for neuroblastoma to assess the inhibitory effect of TDG using different cell lines. Second, in our study, we mainly relied on experimental assays that serve as functional reporters of the progenitor activity of neuroblastoma cell lines, as well as the differentiation and self-renewal ability of the stem/progenitor cell population. Future studies will aim at evaluating the inhibitory effect of TDG, at a molecular level, on different GSK-3β-related signaling pathways that are entangled in the pathophysiology of neuroblastoma and its CSCs. Based on future experiments and knowing that TDG has made it into clinical trials for Alzheimer’s disease, it is thus worthy of consideration in nervous system tumors as well such as neuroblastoma, especially that we proved in our current study its efficiency in targeting the CSC population in this tumor type.

## 5. Conclusions

In conclusion, TDG proved to be an effective *in vitro* and *in vivo* treatment for neuroblastoma cell lines and may hence serve as a potential adjuvant therapeutic agent for this aggressive nervous system tumor. Henceforth, our study supports the notion that targeting GSK-3β, causing decrease in CSCs viability, may be crucial to halt neuroblastoma tumor progression.

## 6. Acknowledgments

We would like to thank all members in the Abou-Kheir’s Laboratory (The WAK Lab) for their help on this work. In addition, we would like to thank members of the core facilities in the DTS Building at the American University of Beirut (AUB) for their help and support.

## 7. Financial Disclosure

This work was supported by the Lebanese National Counsil for Scientific Research Grant Research Program (LNCSR-GRP) (to YF) and the Neuroscience Research Center, Faculty of Medicine, Lebanese University (LU) (to HH). Funders had no role in the study design; in the collection, analysis and interpretation of data; in the writing of the report; and in the decision to submit the article for publication.

## Supplementary Figures

**Supp Fig 1.**
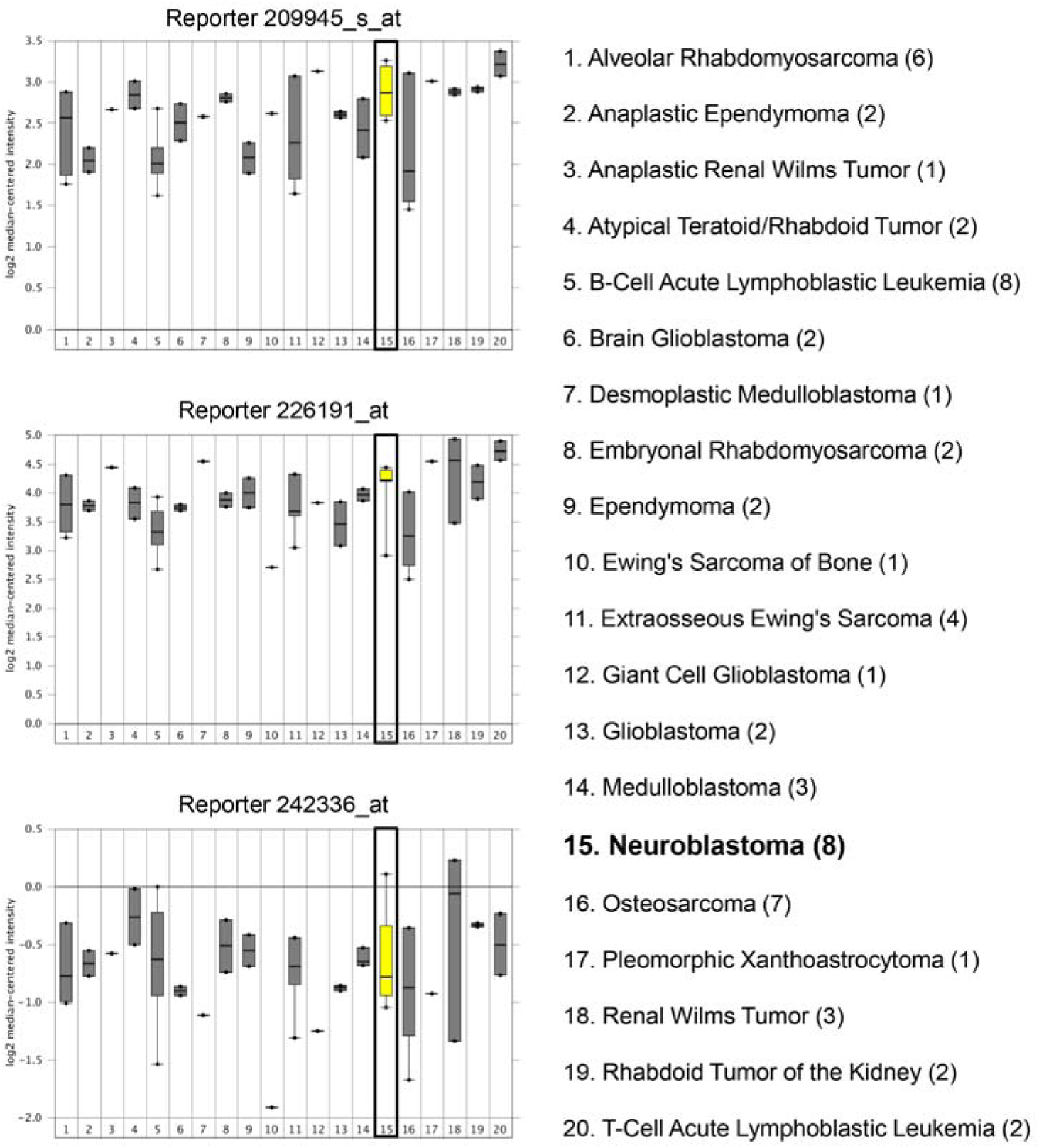
Assessment of the expression levels of *GSK-3β* mRNA in three out of four probes of the Neale Multi-cancer Statistics array set comprised of human pan-tumor samples. Data from different probes of the same dataset are shown: Reporters 209945_s_at (upper panel), 226191_at (middle panel), and 242336_at (lower panel). Expression within tumor tissues was presented by log (base 2) median-centered expression of *GSK-3β*. Box and whiskers plots indicate median and interquartile range. *p*-values were obtained using *t*-tests (Neale Multi-cancer Statistics, 60 samples; data retrieved from Oncomine.org). Analysis revealed that mRNA expression of *GSK-3β* gene was amongst the highest in neuroblastoma tissues relative to other organ specific tumor tissues in reporters 209945_s_at and 226191_at.

**Supp Fig 2.**
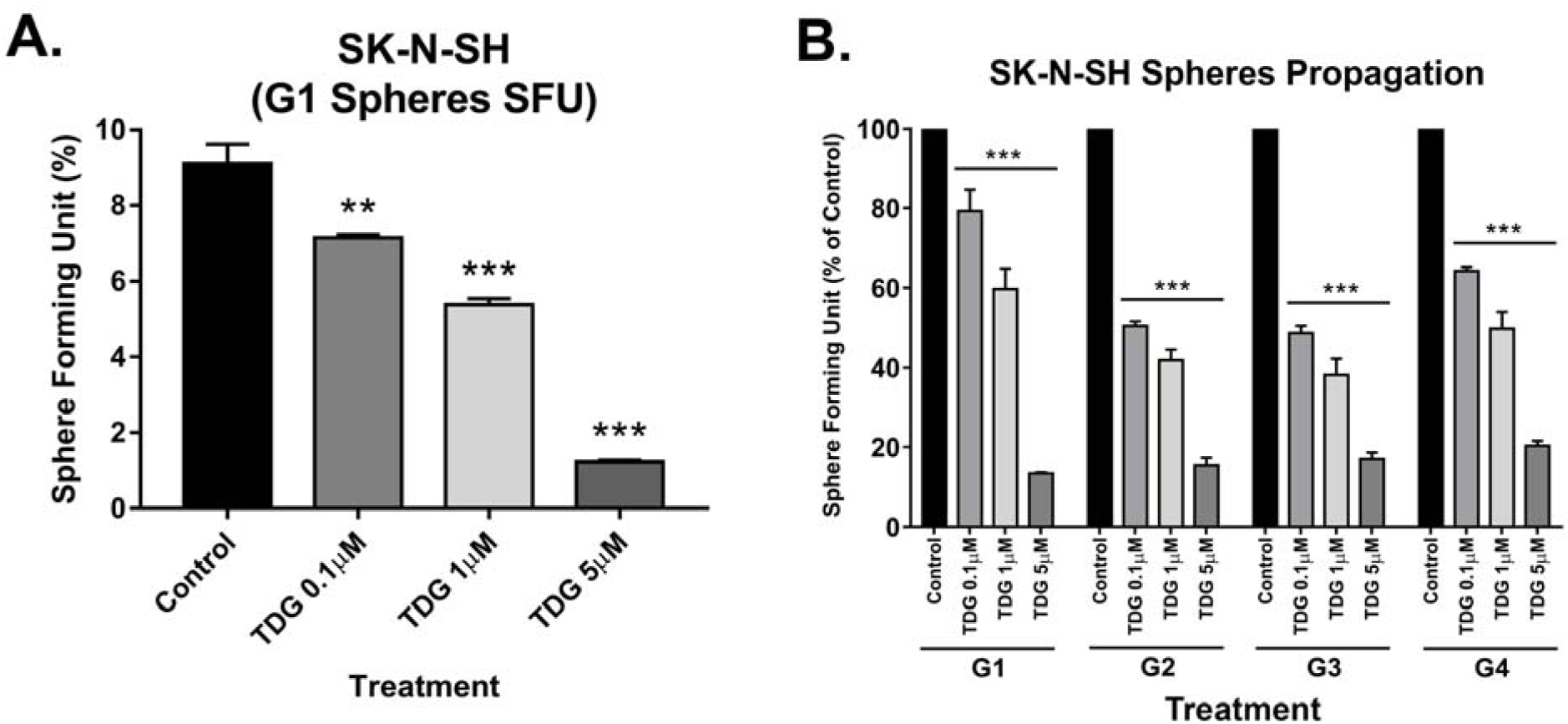
Tideglusib effectively decreases the percentage of SFUs of spheres in the sphere formation assay on SK-N-SH cells and targets an enriched cancer stem/progenitor subpopulation within those cells, decreasing SFU across multiple generations. **(A)** Tideglusib decreases the percentage of SFUs in SK-N-SH cell suspensions in a dose-dependent manner. (One-way ANOVA followed by Bonferroni multiple comparisons: treatment F_(3, 8)_ = 147.6, *p<0*.*001)*. **(B)** SFU obtained from serially passaged SK-N-SH spheres over 4 generations (G1□G4) is shown under untreated condition (control) and with increasing concentration of TDG: 0.1, 1, and 5μM (treated at each generation from G1 to G4) (two-way ANOVA with repeated measures: treatment F_(3, 32)_ = 18.92, *p<0*.*001*; generation F_(3, 32)_ = 688.6, *p<0*.*001*; interaction F_(9, 32)_ = 8.084, *p<0*.*001*). *The data are reported as mean ± SEM of three independent experiments. Bonferroni post-hoc analysis was done to determine simple factor effects. (**P<0*.*01 and ***p<0*.*001 when compared to control)*.

**Supp Fig 3.**
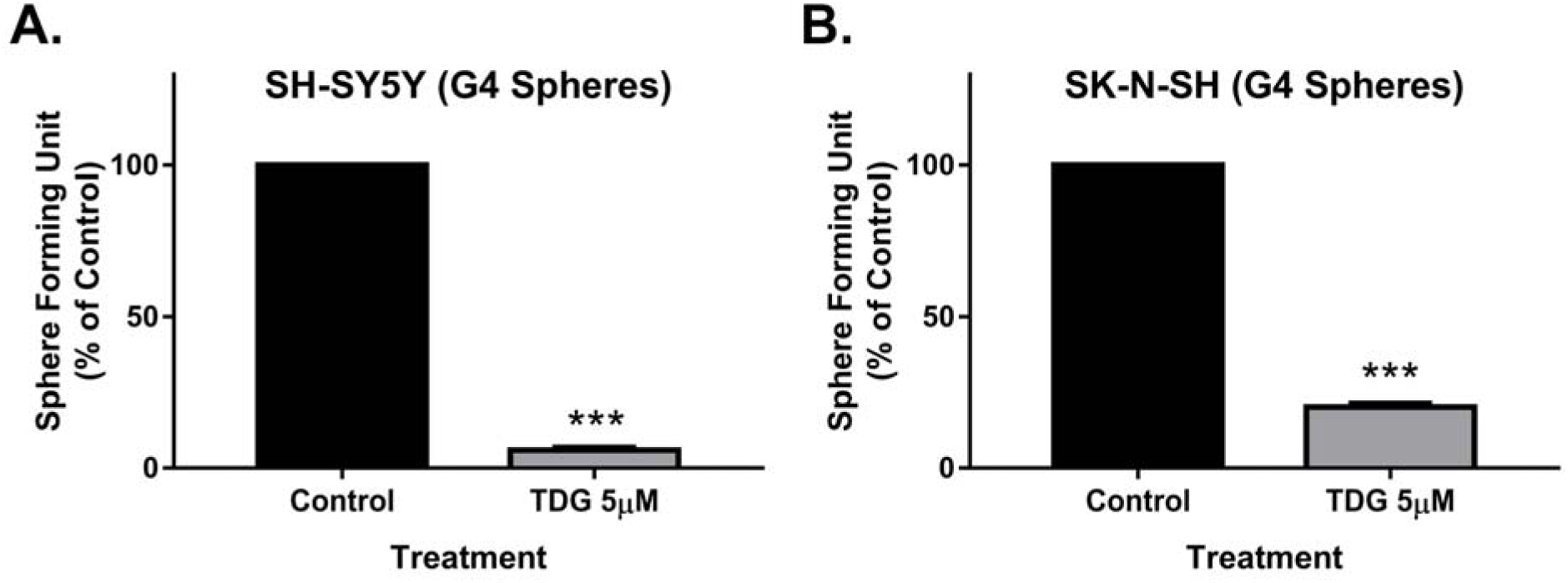
Treating an enriched cancer stem/progenitor subpopulation within SH-SY5Y and SK-N-SH cells significantly decreased SFU at G4. Treating an enriched cancer stem/progenitor subpopulation in SH-SY5Y **(A)** and SK-N-SH **(B)** cells leads to a significant decrease in the percentage of SFUs at G4. *(***P<0*.*001; treatment compared to control, student independent t-test)*.

**Supp Fig 4.**
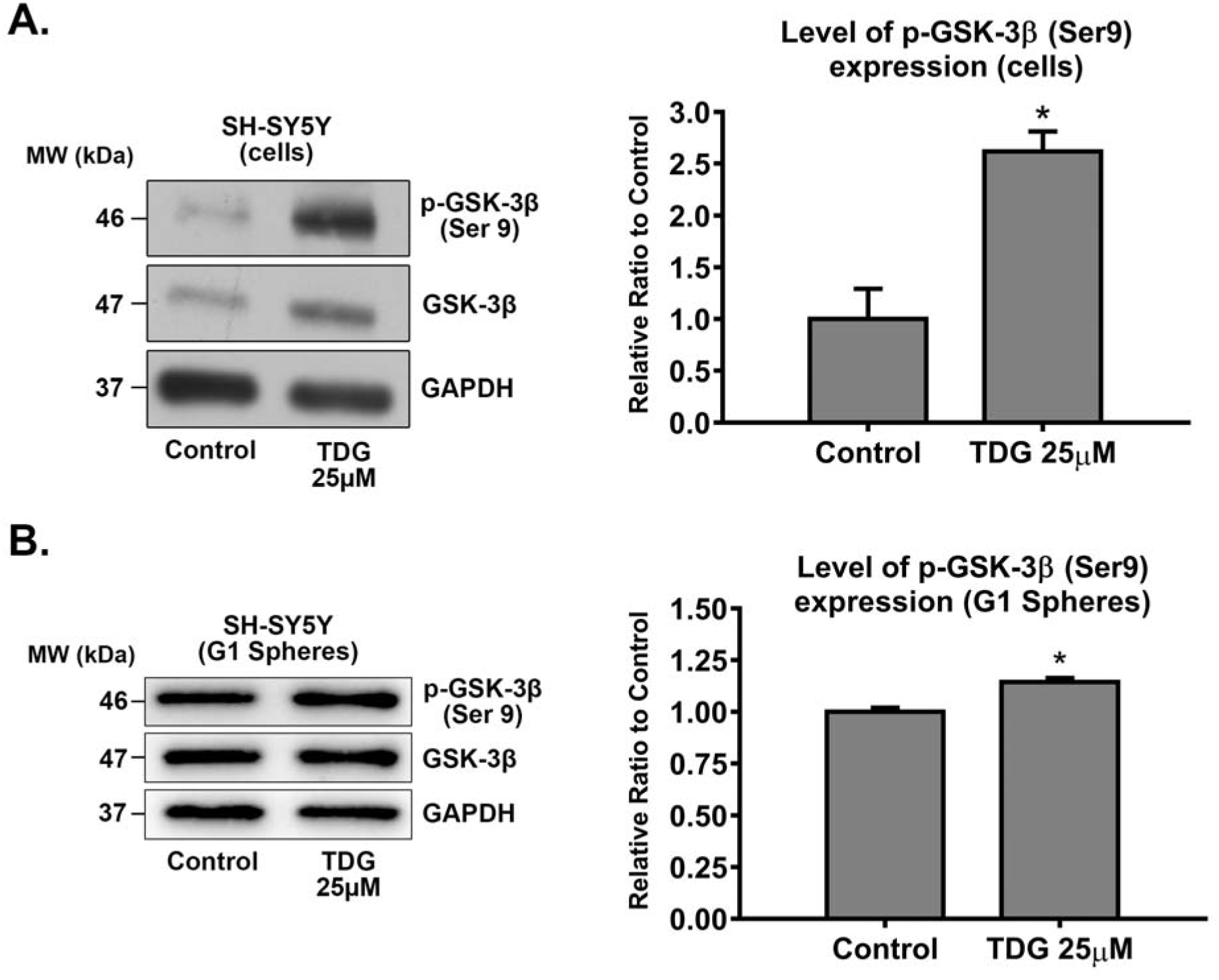
Tideglusib selectively inhibits GSK-3β by increasing expression of its inhibited form, phosphorylated at Serine 9 (p-GSK-3β Ser 9). After treating SH-SY5Y cells with 25μM TDG (for 48 hours) **(A)** and G1 spheres with 5μM TDG **(B)**, proteins were extracted using RIPA buffer, and used to detect differences in expression of the phosphorylated form of GSK-3β (Ser 9). Bands were detected by enhanced chemiluminescence (ECL) using ChemiDoc MP Imaging System. Protein expression was quantified using Image Lab software, relative to the expression of GAPDH, a housekeeping gene equally expressed in treated and non-treated cells/spheres. Results are expressed as relative ratio to control. Analysis of p-GSK-3β (Ser 9) protein level was done after normalization with total GSK-3β protein levels. Data represent an average of three independent experiments. The data are reported as mean ± SEM. *(*P<0*.*05; treatment compared to control, student independent t-test)*.

